# Enhancer RNA like function of intergenic inherited lncRNAs during maternal to zygotic transition in zebrafish

**DOI:** 10.64898/2026.08.15.744761

**Authors:** Dheeraj Chandra Joshi, Sauranil Guha, Nabeel Ahmed, Sanovar Dayal, Beena Pillai

## Abstract

The maternal-to-zygotic transition (MZT) is a major developmental event during which inherited transcripts are remodeled and zygotic transcription is established. Although parentally inherited long noncoding RNAs (lncRNAs) are present in early embryos, they have been thought to be dispensable. We have identified more than 2000 inherited lncRNAs in zebrafish embryos, but how these RNAs participate in regulatory programs during early development has remained unexplored. Here, the inheritance of selected zebrafish lncRNAs spanning a broad expression range were confirmed at the pre-MZT stage and full-length sequences were captured by Direct RNA nanopore sequencing. We show that 30% inherited intergenic lncRNAs are preferentially associated with active enhancers, annotated as such in DANIO CODE, whereas non-inherited intergenic lncRNAs rarely overlap with enhancers. Perturbation of five inherited intergenic lncRNAs, individually, using antisense oligonucleotides reduced the expression of their respective neighboring genes at 2.5, 4.3, and/or 6 hours post fertilization, indicating that these RNAs act as positive local regulators during MZT. Together, these findings identify inherited intergenic lncRNAs as enhancer-associated regulators with elncRNA-like properties during early embryogenesis.

## Introduction

Fertilization initiates a developmental program in which transcriptionally silent gametes give rise to a totipotent embryo. In the earliest stages, embryogenesis is directed primarily by inherited factors, and this control is progressively transferred to the zygotic genome during the maternal-to-zygotic transition (MZT) (Jukam, Shariati, and Skotheim 2017; Schulz and Harrison 2019). In zebrafish, this transition occurs around 3 hours post fertilization (hpf), providing a well-defined and experimentally accessible system for investigating how inherited factors shape early gene-regulatory programs (Harvey et al. 2013; M. T. Lee, Bonneau, and Giraldez 2014).

The contribution of inherited mRNAs and proteins to early development has been extensively studied, but the functions of inherited long noncoding RNAs (lncRNAs) were found to be dispensable for embryogenesis, viability and fertility (Goudarzi et al. 2019). lncRNAs can regulate gene expression through diverse transcriptional, chromatin-associated, and post-transcriptional mechanisms, acting either locally in cis or at distant genomic loci in trans (Yan et al. 2017; Statello et al. 2021). Cis-acting lncRNAs are defined by their ability to regulate genes in a manner dependent on, or linked to, their site of transcription. They can activate, repress, or otherwise modulate nearby gene expression through the RNA product, the act of transcription, local DNA regulatory elements, or combinations of these features (Gil and Ulitsky 2020). Importantly, local proximity alone does not establish cis-regulation, making targeted perturbation essential for assigning this mode of action (Dhaka et al. 2024). Our previous work demonstrated inheritance of the conserved lncRNA Cyrano in zebrafish embryos (Mayuresh Anant Sarangdhar et al. 2018), a lncRNA previously implicated in zebrafish brain development (Ulitsky et al. 2011). Multiple studies demonstrate that lncRNAs can regulate nearby genes in cis through diverse activating or repressive mechanisms (Ponjavic et al. 2009; Werner and Ruthenburg 2015; Quinodoz et al. 2021; Brown et al. 1991; Chen et al. 2014; Wang et al. 2011; Luo et al. 2016). We also showed that lncRNAs transcribed from the kalrn locus, including zebrafish durga and mouse Kalnc2, positively regulate expression of the nearby kalrn gene (Mayuresh A. Sarangdhar et al. 2017; Pal et al. 2023). These findings support the possibility that local lncRNA-mediated regulation is relevant in vertebrate development and provide a rationale for investigating inherited lncRNAs as potential regulators of neighboring genes during MZT.

Previous work from our group identified approximately 2000 inherited lncRNAs in zebrafish by meta-analysis of publicly available RNA-sequencing datasets (Joshi et al. 2025). Here, we experimentally validate a subset of inherited lncRNAs using RT-PCR and Direct RNA nanopore sequencing, demonstrating the presence of intact, full-length transcripts in 2-cell embryos before initiation of zygotic transcription. We then show that inherited intergenic lncRNAs are significantly enriched for overlap with active enhancer regions across developmental stages, whereas non-inherited intergenic lncRNAs are underrepresented. Finally, antisense oligonucleotide-mediated knockdown of selected inherited intergenic lncRNAs reduces expression of neighboring genes during MZT. Together, these findings identify inherited intergenic lncRNAs as an enhancer-associated class of candidate local regulators during early zebrafish development.

## Results

### Inherited lncRNAs show a wide range of expression in 2-cell embryos

To validate inherited lncRNAs experimentally, 14 candidates were shortlisted from the previously defined inherited lncRNA set so as to span a broad expression range and represent both genic and intergenic classes (Table 1, Figure 1A). For ease of reference, these candidates were numbered sequentially with the prefix ilnc, for inherited long non-coding. RT-PCR performed on RNA isolated from 2-cell embryos confirmed the presence of 11 of 14 tested lncRNAs, indicating that inherited lncRNAs are readily detectable in the early embryo across a wide abundance range (Figure 1B).

**Table 1.**
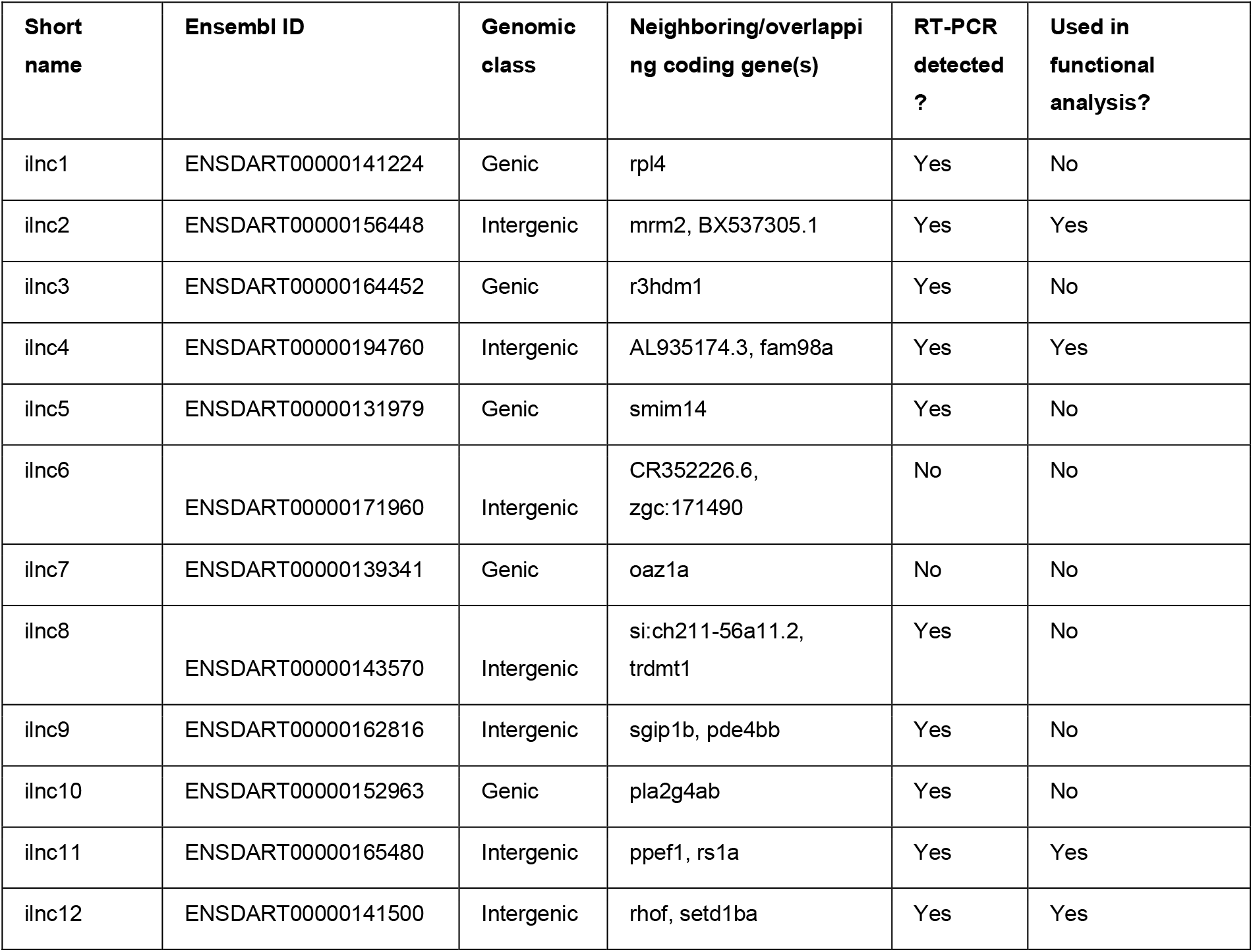
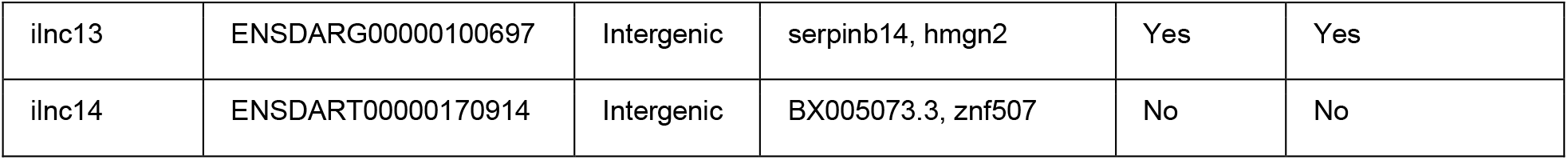
Table summarizing the genomic class and neighboring or overlapping coding genes of the shortlisted inherited lncRNAs.

**Figure 1.**
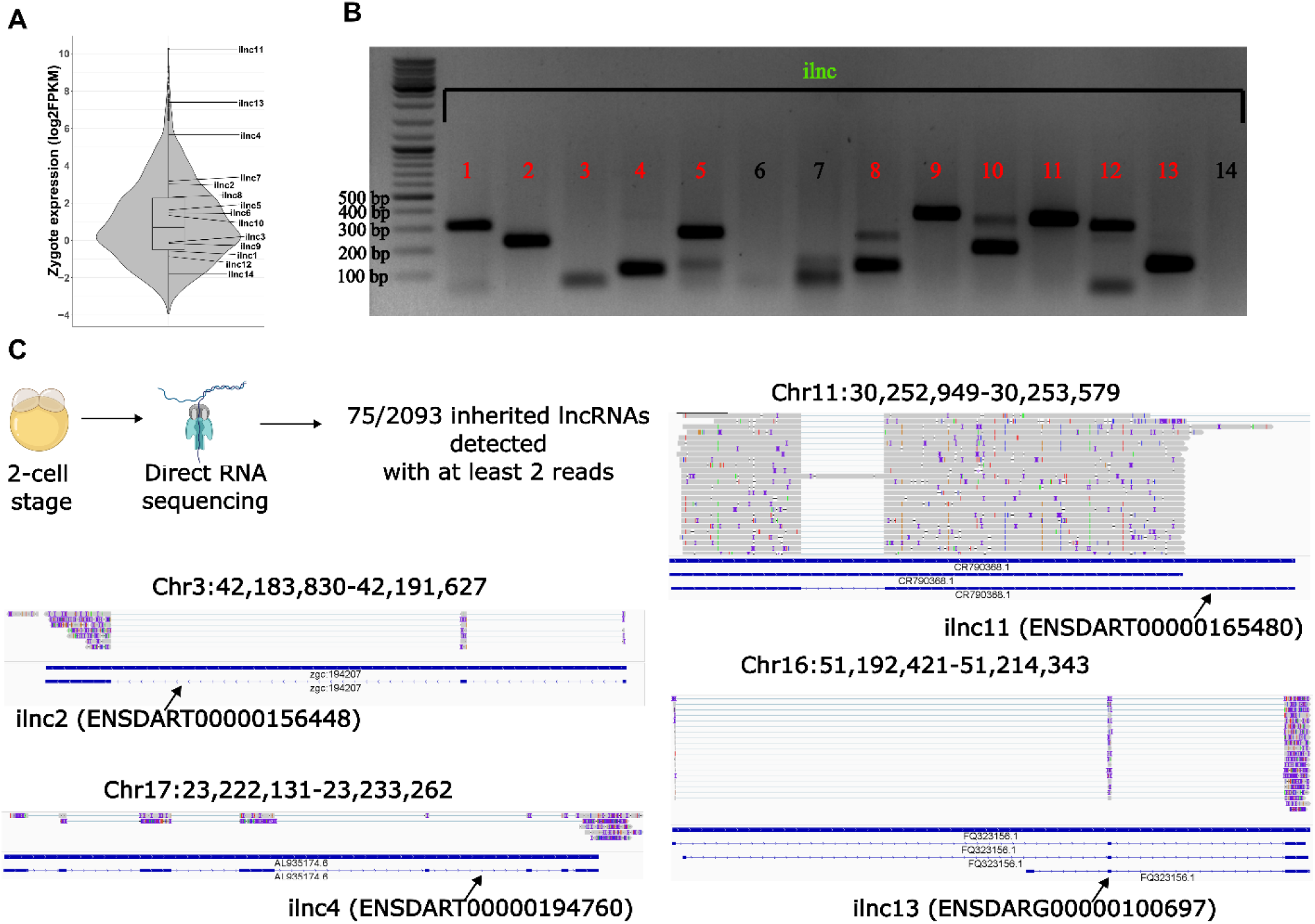
Inherited lncRNAs across the expression range are readily detected by RT-PCR and full-length reads. (A) Overview of the inherited lncRNAs shortlisted for experimental validation, spanning a broad expression range. (B) Endpoint RT-PCR validation of inherited lncRNAs using RNA isolated from 2-cell zebrafish embryos. The majority of shortlisted inherited lncRNAs were detected across the expression range. (C) Integrated Genomics Viewer (IGV) views for representative inherited lncRNA loci (ilnc2, ilnc4, ilnc11, and ilnc13) showing full-length or near full-length read support in 2-cell embryos.

A key unresolved question was whether inherited lncRNAs are present in embryos as intact transcripts or only as fragmented molecules. To address this, direct RNA nanopore sequencing was performed on RNA isolated from 2-cell embryos. Using a minimum detection threshold of two reads per transcript, 75 inherited lncRNAs were detected in the nanopore dataset. Among the experimentally prioritized loci, full-length or near full-length read support was obtained for four inherited lncRNAs: ilnc2, ilnc4, ilnc11, and ilnc13 (Figure 1C). These data strengthen the conclusion that complete inherited lncRNA transcripts are present in the early embryo. Minor differences between nanopore-supported transcript termini and existing but shorter Ensembl annotations were observed for ilnc4 and ilnc11, suggesting that inherited isoforms at these loci may be incompletely annotated. Additionally, the presence of inherited lncRNAs was also confirmed in the reanalysis of publicly available long-read transcriptomics dataset of fertilized zebrafish zygote (Supplementary figure 1). Overall, this validation step provided orthogonal confirmation of inherited lncRNA presence beyond the previous short-read meta-analysis. Also, it established a practical subset of experimentally tractable and fully verified loci for downstream analysis.

### Inherited intergenic lncRNAs preferentially overlap active enhancers across development

To place inherited lncRNAs in a broader regulatory context, active enhancer annotations from DANIO-CODE were intersected with zebrafish lncRNA coordinates (Baranasic et al. 2022). The analysis was stratified into inherited versus non-inherited and genic versus intergenic lncRNAs (Figure 2A).

**Figure 2.**
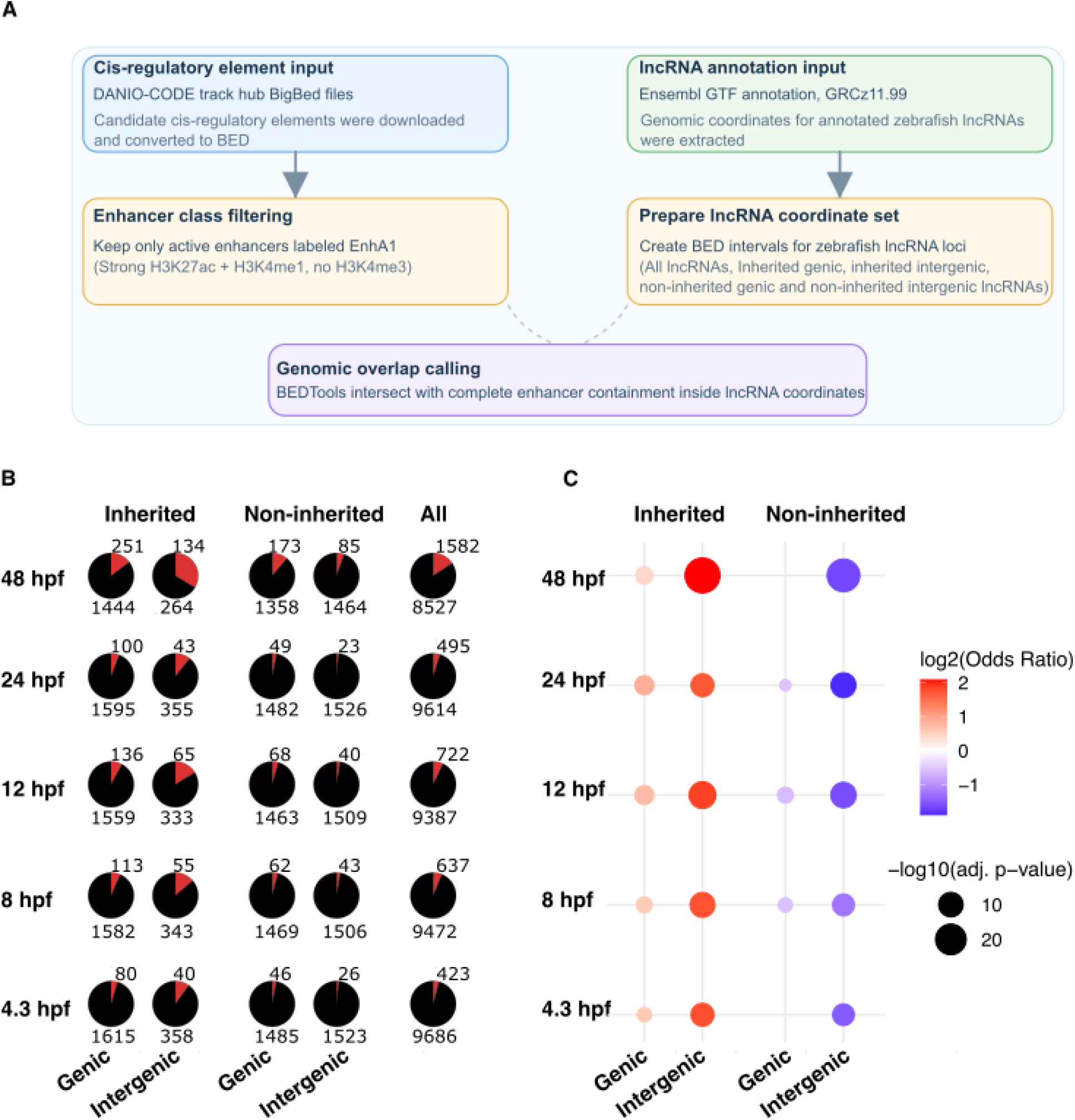
Intergenic inherited lncRNAs preferentially overlap with enhancer elements across development. (A) Schematic of the computational workflow used to identify enhancer-overlapping lncRNAs. (B) Pie charts showing the proportion of enhancer-overlapping (red) versus non-overlapping (black) lncRNAs at 4.3, 8, 12, 24, and 48 hpf, stratified into inherited genic, inherited intergenic, non-inherited genic, non-inherited intergenic, and all lncRNAs. (C) Enrichment analysis (odds ratio and Fisher’s exact test) summarizing the association between enhancer overlap and lncRNA class across developmental stages.

Across all developmental stages examined, inherited intergenic lncRNAs were significantly enriched for overlap with active enhancers (Figure 2B and 2C). This enrichment was strongest at 48 hpf, where inherited intergenic loci showed an odds ratio of 4.25 (134 overlapping loci; adjusted *p* = 6.77 × 10^−30^) and remained robust at 24 hpf (odds ratio 3.24; adjusted *p* = 7.03 × 10^−9^), 12 hpf (odds ratio 3.62; adjusted *p* = 6.11 × 10^−14^), 8 hpf (odds ratio 3.35; adjusted *p* = 3.10 × 10^−11^), and 4.3 hpf (odds ratio 3.40; adjusted *p* = 9.03 × 10^−9^). By contrast, non-inherited intergenic lncRNAs were consistently underrepresented among enhancer loci at all stages, with odds ratios ranging from 0.27 to 0.42 and all adjusted p-values significant. Inherited genic lncRNAs also displayed significant enrichment for enhancer overlap, but with substantially smaller effect sizes than the intergenic inherited class, with odds ratios between 1.37 and 1.83. These findings identify inherited intergenic lncRNAs as the clearest enhancer-associated subclass within the inherited lncRNA pool and suggest that they constitute a distinct group of candidate enhancer-associated local regulators during embryogenesis.

### Inherited intergenic lncRNAs positively regulate neighboring genes during MZT

The enhancer overlap analysis suggested that inherited intergenic lncRNAs may function as local positive regulators. To test this directly, antisense oligonucleotide (ASO)-mediated knockdown experiments were conducted on the five intergenic inherited lncRNAs examined functionally: ilnc2, ilnc4, ilnc11, ilnc12, and ilnc13 (Figure 3A).

**Figure 3.**
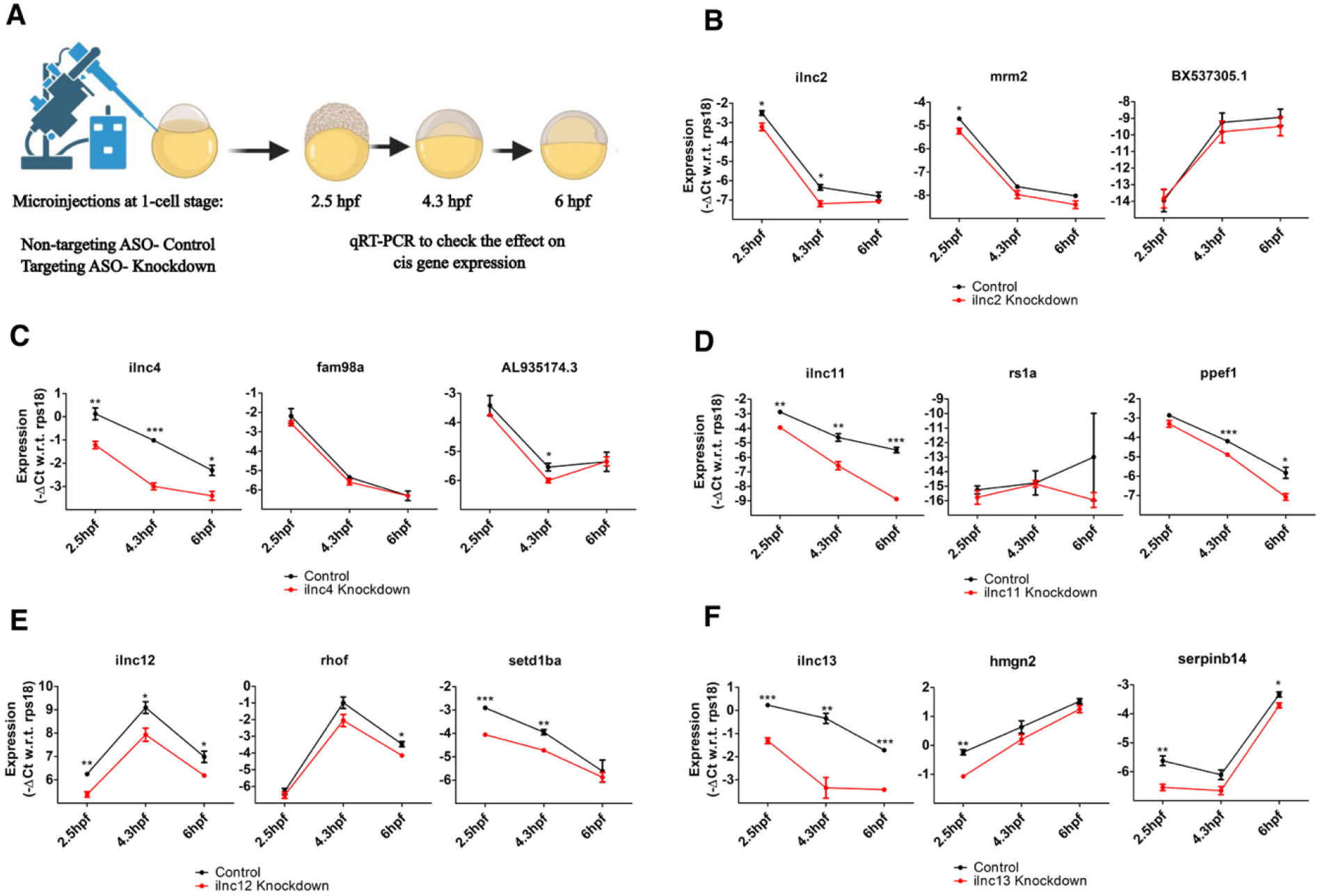
Inherited intergenic lncRNAs positively regulate neighboring genes during the maternal-to-zygotic transition. (A) Experimental outline for ASO-mediated knockdown of inherited intergenic lncRNAs followed by RT-qPCR analysis of neighboring gene expression at 2.5, 4.3, and 6 hpf. (B-F) Relative expression analysis of neighboring genes after knockdown of ilnc2, ilnc4, ilnc11, ilnc12, and ilnc13, respectively. At least 3 biological replicates were taken for every condition. Statistical significance was determined using a two-tailed Student’s t-test, *p < 0.05, **p < 0.01, ***p < 0.001.

Knockdown of ilnc2 reduced expression of its neighboring gene mrm2 at 2.5 hpf, with expression tending to recover at later stages (Figure 3B). Knockdown of ilnc4 resulted in transient downregulation of AL935174.3 at 4.3 hpf (Figure 3C). Knockdown of ilnc11 reduced expression of ppef1 at 4.3 hpf and 6 hpf (Figure 3D). Knockdown of ilnc12 reduced setd1ba at 2.5 hpf and 4.3 hpf, while rhof was reduced at 6 hpf (Figure 3E). Finally, knockdown of ilnc13 reduced hmgn2 at 2.5 hpf and serpinb14 at 2.5 hpf and 6 hpf (Figure 3F). Taken together, these results show that inherited intergenic lncRNAs positively regulate expression of neighboring genes during MZT.

Importantly, the time-resolved effects differed among loci. Some targets were affected at pre-ZGA stages, whereas others were altered predominantly after zygotic transcription had begun. This stage-specificity suggests that inherited intergenic lncRNAs may not operate through a single mechanism at all loci. Instead, they may influence local gene output through distinct enhancer-associated or post-transcriptional processes depending on locus context and developmental stage.

Embryos injected with ASOs targeting inherited intergenic lncRNAs were also followed through early larval stages and examined for overt morphological abnormalities at 4 dpf. Under the conditions tested, no obvious differences in gross morphology or survival were observed between control and knockdown embryos (Supplementary figure 2). The absence of overt phenotypes indicates that the early expression effects reported here are lineage-restricted, rather than globally disruptive to embryogenesis. Interestingly, ilnc1, ilnc2, ilnc4, and ilnc12 have syntenically conserved counterparts in both mouse and human inherited RNA pools, raising the possibility of similar positive cis-regulation in mammals (Supplementary figure 3)(Joshi et al. 2025).

## Materials and Methods

### Animal husbandry

Zebrafish (Danio rerio) of the Assam wild-type (ASWT) strain were maintained at 28 °C under a 14 h light/10 h dark cycle in the zebrafish facility at CSIR–Institute of Genomics and Integrative Biology (CSIR-IGIB), following standard zebrafish husbandry procedures. Male and female fish were paired in mating tanks overnight, and fertilized embryos were collected the following morning for microinjection experiments. All animal handling and experiments were conducted in accordance with the guidelines approved by the Institutional Animal Ethics Committee (IAEC) of CSIR-IGIB, India.

### Candidate inherited lncRNA selection

Inherited lncRNAs were selected from the previously defined zebrafish inherited lncRNA dataset (Joshi et al. 2025). Fourteen candidates were shortlisted to represent a broad expression range and both genic and intergenic classes. For ease of presentation, these loci were renamed sequentially as ilnc1 to ilnc14.

### Identification of enhancer-overlapping lncRNAs

To identify enhancer-associated inherited lncRNAs, BigBed files containing cis-regulatory element annotations were obtained from the DANIO-CODE track hub (Baranasic et al. 2022). BigBed files were converted to BED format using the UCSC bigBedToBed utility. Only active enhancers annotated as EnhA1 were retained for downstream analysis; this class corresponds to regions marked by strong H3K27ac and H3K4me1 and lacking H3K4me3.

Zebrafish lncRNA genomic coordinates were extracted from the Ensembl GTF annotation (GRCz11.99). BED intervals were generated for all annotated lncRNAs and then partitioned into inherited genic, inherited intergenic, non-inherited genic, and non-inherited intergenic sets. Genomic overlaps between lncRNA loci and enhancer intervals were identified using BEDTools intersect with the -f 1.0 option, requiring complete overlap of enhancer coordinates within lncRNA loci (Quinlan 2014). The number of unique enhancer-overlapping lncRNAs was quantified using standard UNIX command-line tools, and the results were visualized in R (v4.5.1) using custom plotting functions.

### RNA isolation from zebrafish embryos

Embryos were collected at the indicated developmental stages (2-cell, 2.5, 4.3, and 6 hpf) in RNAiso Plus (Takara, 9109) and stored at ^−^20 °C until processing. RNA was isolated from 20 embryos for RT-PCR validation, 25 embryos for nanopore sequencing, and 3–15 embryos for RT-qPCR. Embryos were homogenized in RNAiso Plus, followed by chloroform extraction and isopropanol precipitation. RNA pellets were washed with 70% ethanol prepared in 0.1% DEPC-treated water, air-dried briefly, and resuspended in nuclease-free water. RNA concentration was measured using a NanoDrop spectrophotometer, and purified RNA was stored at ^−^20 °C until further use.

### cDNA synthesis and RT-PCR

For endpoint RT-PCR, 500 ng total RNA was treated with DNase I (Ambion, AM2222) and reverse-transcribed using random primers (Promega, C1181) and M-MuLV Reverse Transcriptase (NEB, M0253). lncRNA-specific primers are listed in Supplementary Table 1. PCR was performed using KAPA SYBR FAST chemistry (Merck, KK4618), and products were resolved on 1.5% agarose gels.

### Direct RNA nanopore sequencing

Total RNA from 25 2-cell-stage embryos was used for direct RNA sequencing. Libraries were prepared from 500 ng total RNA using the Oxford Nanopore Technologies Direct RNA Sequencing Kit (SQK-RNA002) and sequenced on an R9.4.1 flow cell (FLO-MIN106) using a MinION Mk1C device. Reads were aligned to the zebrafish GRCz11 reference transcriptome using minimap2, and transcript abundance was quantified using Salmon v1.10.3. Splice-aware minimap2 alignments were visualized in IGV to assess full-length reads mapping to inherited lncRNAs.

### Publicly available PacBio sequencing analysis

Fastq.gz format was downloaded for the run id SRR32588735 (0 hpf) from NCBI. Long-read sequencing compatible genome alignment was performed with minimap2 in spliced-alignment mode (-ax splice) with an overall mapping rate of 99.93% using GRCz11. Transcripts assembled and quantified using StringTie V3 through Danio_rerio.GRCz11.113.chr.gtf. Violin plots generated using Rstudio. Full-length reads were visualized in IGV.

### ASO microinjections

Fertilized embryos were collected 15–20 min after spawning and injected at the one-cell stage with 2 nL of lncRNA-targeting or non-targeting control ASOs (50 ng/µL; 100 pg per embryo). Injections were performed using fine glass capillaries with an Eppendorf FemtoJet 4i microinjector mounted on a Nikon SMZ800N stereomicroscope. Injected embryos were maintained at 28 °C until collection at the indicated stages.

### cDNA synthesis and RT-qPCR

For RT-qPCR, 500 ng total RNA was treated with DNase I and reverse-transcribed using random primers and M-MuLV Reverse Transcriptase. No-reverse-transcriptase controls were included for each sample. RT-qPCR was performed in 10 µL reactions using KAPA SYBR FAST chemistry and gene-specific primers in 384-well plates. Each reaction was run in three technical replicates, alongside no-template controls. Amplification was performed for 40 cycles using the following conditions: 95 °C for 15 s, 60 °C for 15 s, and 72 °C for 20 s, following an initial denaturation step at 95 °C for 3 min.

### Morphological imaging

Larvae were examined at 4 dpf for gross morphological abnormalities. Before imaging, larvae were immobilized in methylcellulose, and brightfield images were acquired using a Nikon SMZ800N stereomicroscope.

## Discussion

In this study, we identify inherited intergenic lncRNAs as a distinct class of candidate local regulators during the zebrafish maternal-to-zygotic transition. RT-PCR validation and Direct RNA nanopore sequencing confirmed that inherited lncRNAs, as full-length transcripts, are present in early embryos. Importantly, inherited intergenic lncRNAs were significantly enriched for overlap with active enhancer elements across all developmental stages examined, with odds ratios ranging from 3.24 to 4.25. In contrast, non-inherited intergenic lncRNAs were consistently depleted for enhancer overlap, with odds ratios between 0.27 and 0.42. Together with the weaker enrichment observed for inherited genic lncRNAs, these results identify inherited intergenic lncRNAs as the most strongly enhancer-associated subclass within the inherited lncRNA pool.

The enhancer association provides a genome-wide framework for functional perturbation experiments. Knockdown of five inherited intergenic lncRNAs reduced expression of one or more neighboring genes during MZT, supporting a role in positive local regulation. This behavior is consistent with studies of enhancer-associated lncRNAs or elncRNAs, which can be linked to activation of nearby genes, enhancer activity, and local chromatin interactions (Ørom et al. 2010; Ørom and Shiekhattar 2013; Tan et al. 2020). The combination of active-enhancer overlap and target-gene downregulation following lncRNA depletion therefore shows that inherited intergenic lncRNAs are enhancer-associated candidate regulators during early embryogenesis.

However, the present data do not establish that these inherited lncRNAs are bona fide eRNAs or define the precise mechanism underlying their local effects. Enhancer overlap and ASO-mediated knockdown cannot distinguish whether the functional unit is the mature RNA transcript, transcription through the locus, the lncRNA promoter, the underlying enhancer DNA sequence, or a combination of these features. In future, investigations can delineate if these inherited lncRNAs regulate neighboring genes through promoter activity, transcription, or splicing (Engreitz et al. 2016).

The temporal pattern of target-gene effects suggests that different loci may operate through distinct mechanisms. Effects observed at 2.5 hpf, before widespread zygotic genome activation, could reflect post-transcriptional regulation of inherited target transcripts, including altered RNA stability, processing, localization, or translation. In contrast, targets affected predominantly after ZGA, such as ppef1, AL935174.3, and rhof, may be more compatible with transcription-linked local regulation. These possibilities are not mutually exclusive, particularly because maternally inherited transcripts remain a substantial component of the embryonic transcriptome after ZGA (Bhat et al. 2023). Nascent-transcription assays, orthogonal perturbations, enhancer-state profiling, and rescue experiments can distinguish RNA-dependent, transcription-dependent, and DNA-element-dependent functions at individual loci in the future.

Finally, the absence of overt gross morphological defects at 4 dpf does not exclude meaningful developmental functions for inherited intergenic lncRNAs. Transient molecular changes may be buffered by feedback regulation or redundancy, while lineage-specific, physiological, or behavioral phenotypes would not necessarily be detected by brightfield morphology alone. Overall, our findings support a model in which inherited intergenic lncRNAs are selectively associated with active enhancer regions and contribute to positive local gene regulation during MZT. This work establishes these lncRNAs as a tractable class for future mechanistic studies of enhancer-associated regulation in vertebrate embryogenesis.

