## Supplementary Figures and Tables for "Enhancer RNA like function of intergenic inherited lncRNAs during maternal to zygotic transition in zebrafish"

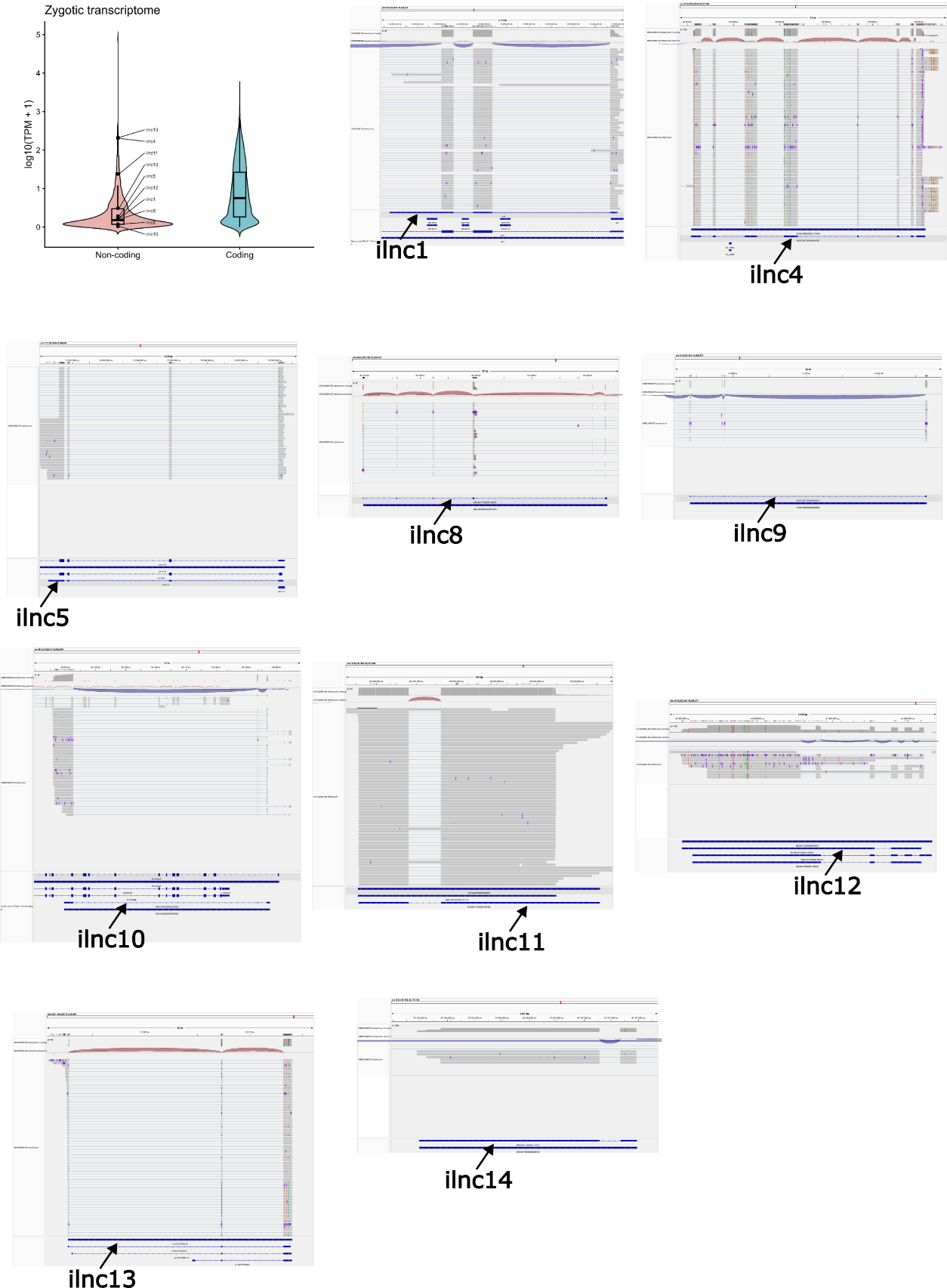

**Supplementary Figure 1. Inherited lncRNAs are detected in publicly available long read sequencing data.** Violin plot showing the detection of shortlisted inherited lncRNAs in a publicly available long-read transcriptomics dataset (SRR32588735) of fertilized zebrafish zygote.

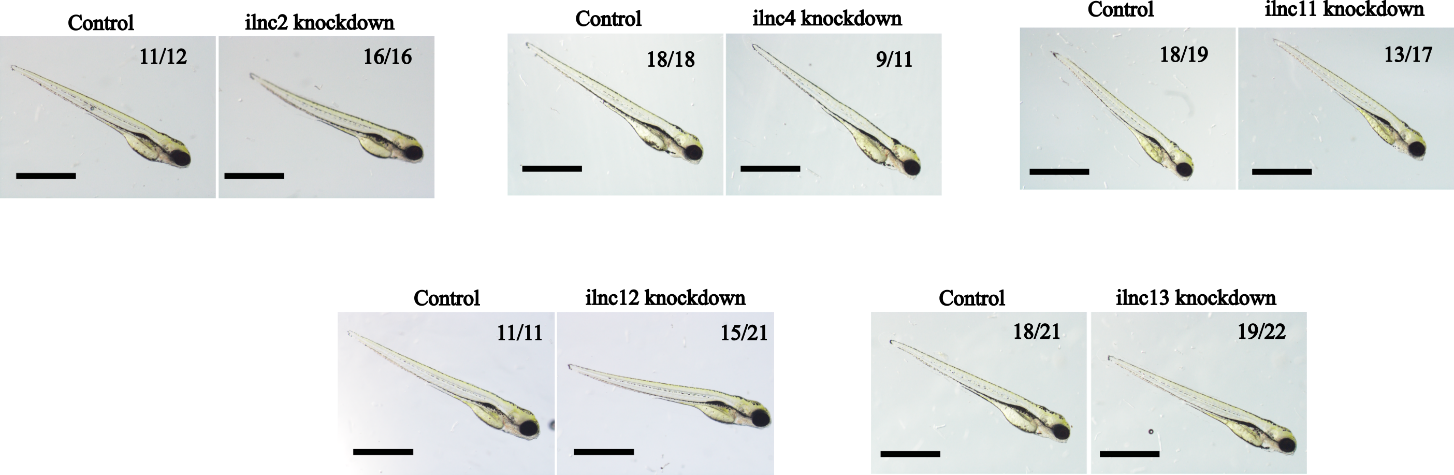

**Supplementary Figure 2. Inherited lncRNA knockdowns have no overt morphological and survival defects.** 4 dpf (days post fertilization) embryos injected with control or ASOs targeting inherited intergenic lncRNAs. The numbers written in the top right corner represent the number of larvae, with morphology shown in the picture out of the total number of embryos scored. Scale bar - 1 mm.

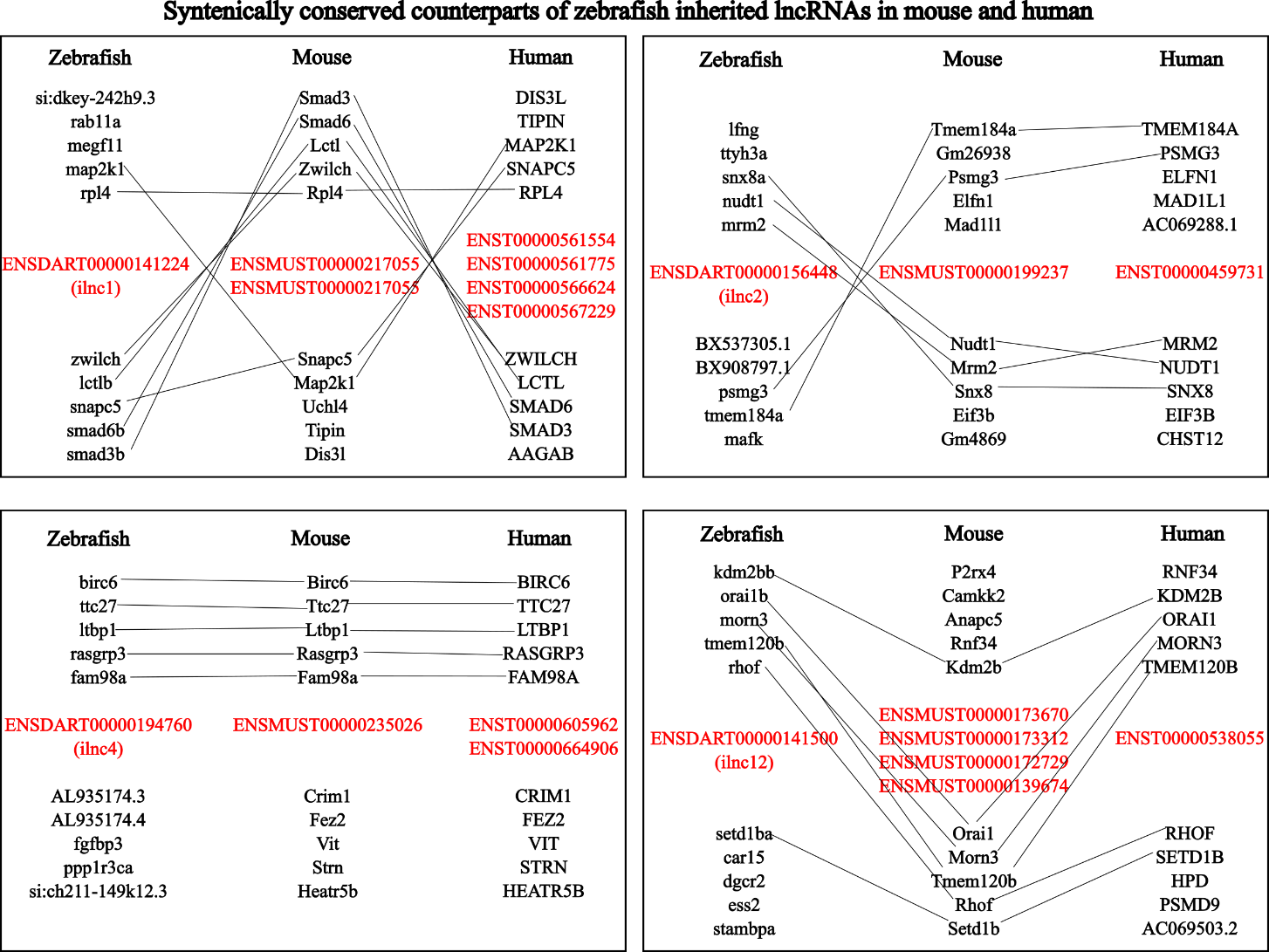

**Supplementary Figure 3. Functionally studied lncRNAs have conserved counterparts in mice and humans.** Schematic representation of the syntenic conservation of ilnc1, ilnc2, ilnc4, and ilnc12 in mouse and human. The Ensembl transcript IDs of inherited lncRNAs of zebrafish, mice, and humans are written in red. The flanking protein coding genes (5 on each side) in the same order as they appear in the chromosomes are written in black. The lines connect the orthologous protein-coding genes in zebrafish, mice, and humans.

**Supplementary Table 1. Primer sequences used for RT-PCR and RT-qPCR.**

| **ilncRNAs** | **Transcript id** | **Forward primer (5’-3’)** | **Reverse primer (5’-3’)** |
| --- | --- | --- | --- |
| ilnc1 | (ENSDART00000141224) | GCTCCCATCCGCCCTGATAT | CCAGCATAAGACAGCAGGCA |
| ilnc2 | (ENSDART00000156448) | GAGCTGTGGTCCCTTCATCC | CAAGCGTTTTGGTGGTGGAA |
| ilnc3 | (ENSDART00000164452) | CATGATCGCACACATCCAGC | CGACTGCGCTTAACATGACC |
| ilnc4 | (ENSDART00000194760) | TCTAGAGGCCAAGCGAGACA | GACCCTCCAGTTCCTTCACC |
| ilnc5 | (ENSDART00000131979) | CACCGAGTGCCTTCAGGACA | TGTGGATGAAATGGAGTTCGATTG |
| ilnc6 | (ENSDART00000171960) | CCCTGTCGGGCCTAACTTTT | GGGCGTCTGAACTAGAGCTG |
| ilnc7 | (ENSDART00000139341) | TATCTGACTGTGCAGCGGTT | AAACGTACACGGCTGTTCGA |
| ilnc8 | (ENSDART00000143570) | ATGAAGCTGGAGAGAGCAGC | CCGCTTGTCCTTGTCCTTGT |
| ilnc9 | (ENSDART00000162816) | GTGTGGATGCTGCTAATGCG | CCCCTCATCATCCACATCAGG |
| ilnc10 | (ENSDART00000152963) | AATGTCTGCTACCGTCGTCA | ACTGTGCTTCTGTTTTCTTGCG |
| ilnc11 | (ENSDART00000165480) | ACAGAGAACACGCCTCATTCA | GCATGCTTCAGGAAAAGAAACG |
| ilnc12 | (ENSDART00000141500) | ACTAGCCACACAACATCGAAC | TGCCTCAAGATTTCCATCAGCT |
| ilnc13 | (ENSDARG00000100697) | TGATCATCAGACTGGGCAGA | GACAGGCTGAGACCCATGAT |
| ilnc14 | (ENSDART00000170914) | GCAAACCTTTCTGCTTGCTT | GGGAGAACGGATACTCGTCA |

| **qRT-PCR primers for** | **Forward primer** | | | **Reverse primer** |
| --- | --- | --- | --- | --- |
| ilnc1 | GCTCCCATCCGCCCTGATAT | | | CCAGCATAAGACAGCAGGCA |
| ilnc13 | TGATCATCAGACTGGGCAGA | | | GACAGGCTGAGACCCATGAT |
| rpl4 | AAGCTCAACCCATACGCCAA | | | TCAGCCTTCTTAGCCTTGGTC |
| hmgn2 | CCAAGATGCCCAAAAGAAAG | | | TGGCCTTTGGTTCTGGTTTA |
| serpinb14 | GTGCGACTCCAGAAGCTCAT | | | GGGCTCCTGGTTTGTTCAGT |
| ilnc2 | GAGCTGTGGTCCCTTCATCC | | | CAAGCGTTTTGGTGGTGGAA |
| ilnc4 | TCTAGAGGCCAAGCGAGACA | | | GACCCTCCAGTTCCTTCACC |
| ilnc11 | ACAGAGAACACGCCTCATTCA | | | GCATGCTTCAGGAAAAGAAACG |
| ilnc12 | ACTAGCCACACAACATCGAAC | | | TGCCTCAAGATTTCCATCAGCT |
| BX537305.1 | TCGCCACAATGCAGAGCTAA | | | CGACTTTGCCGTGCTTTTCA |
| mrm2 | GCTCAGTGACCCGTTCGTTA | | | GCCGTGTGAGTTGAAGGAGA |
| fam98a | CTGTGACTGGTGGTGCTGAA | | | TGGTCAGAGCAGGATACGGA |
| AL935174.3 | GAACAACAGCTTGACGCCTG | | | GGAATAACGCCTCCTCTCCG |
| rs1a | ACATGGGCTGGAAAATCCTG | | | TCCTGCCTCAAAACCCAGAG |
| ppef1 | AGGCACAGATTAAAGTCCAGCA | | | CCGCATCCCATGATCCTACC |
| setd1ba | GGACGGCCATTGACAAAATG | | | GATGGTCTGTATTAGAGACAATGCTG |
| rhof | | TTATACTGATAGGCTGCAAGACG | AGCGTTCATTTCCTTTTGCG | |
| rps18 | | TTCAGCACATCCTTCGTGTC | AGTGAGTTCTCCAGCCCTCTT | |

**Supplementary Table 2. ASO sequences and injection details.**

| **ASOs** | **ASO sequence (* represent phosphorothioate bonds and [mA], [mG], [mC], [mU] - 2'O-Methyl RNA bases)** |
| --- | --- |
| ilnc2 ASO | mU*mC*mG*mA*mA*G*G*G*A*T*C*A*T*T*G*mA*mC*mA*mC*mC |
| ilnc4 ASO | mG*mG*mA*mC*mU*G*A*C*C*A*A*G*C*A*A*mA*mA*mG*mC*mU |
| ilnc11 ASO | mU*mC*mC*mA*mG*A*C*A*G*A*G*C*T*T*G*mC*mU*mU*mC*mG |
| ilnc12 ASO | mG*mC*mA*mC*mA*A*G*G*T*C*A*G*A*T*A*mU*mA*mC*mA*mU |
| ilnc13 ASOs | mU*mG*mA*mC*mG*G*A*G*C*G*G*T*C*A*T*mG*mC*mA*mC*mC and mU*mU*mG*mA*mA*T*A*C*C*T*C*A*G*A*G*mA*mC*mG*mC*mU |
| Non-targeting ASO | mG*mA*mU*mU*mC*C*G*C*G*C*A*C*G*T*G*mA*mG*mU*mA*mG |

**Supplementary Table 3. Enhancer-overlap statistics across developmental stages.**

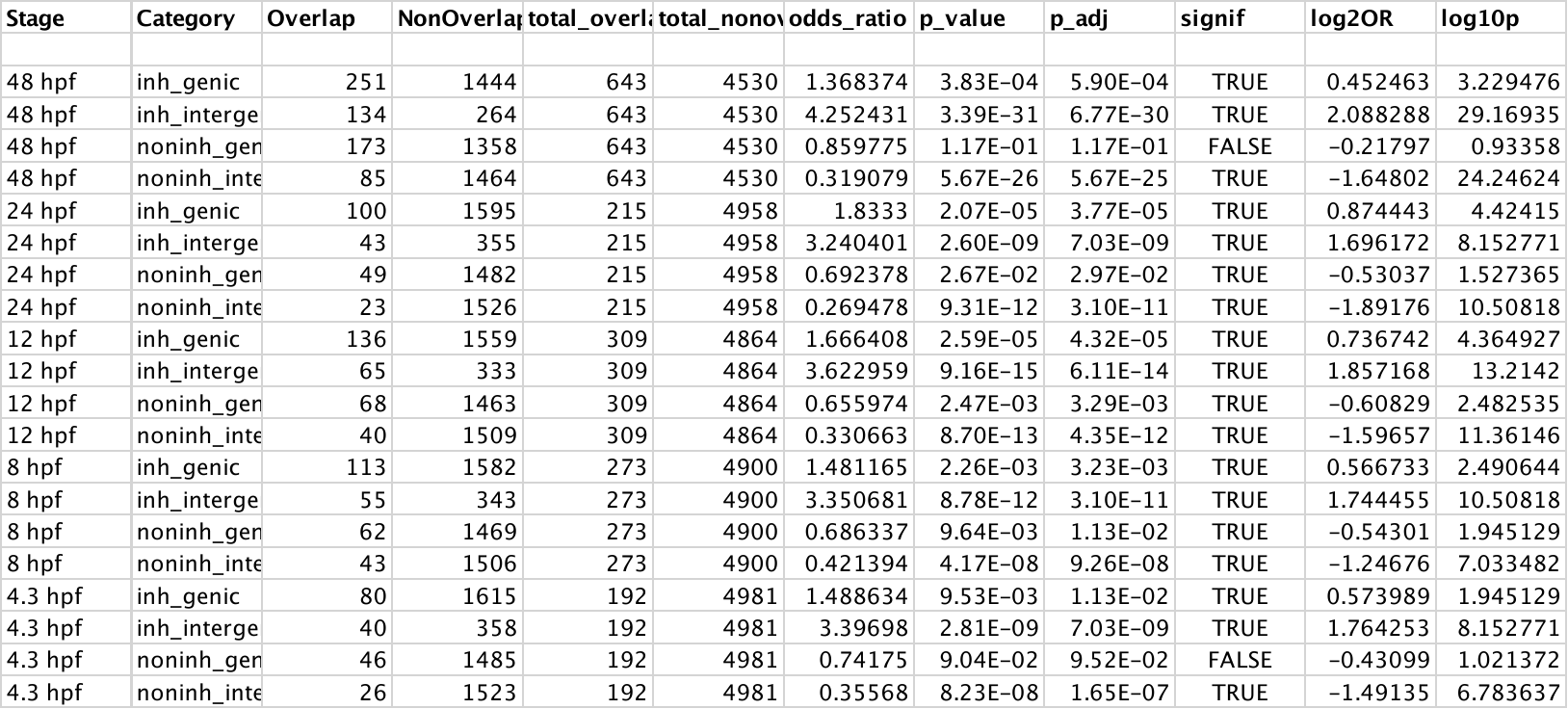
